# Application of MT3 in the Development and Immunogenicity Evaluation of GnRH Vaccines

**DOI:** 10.1101/2025.09.09.675242

**Authors:** Guozhen Zhang, Ying Xu, Yuhan Zhu, Congmei Wu, Yuhe Yin

## Abstract

Research on immune castration vaccines is of great significance for animal management. In this study, five copies of GnRH were linked to two MT3s by a flexible linker, GGGGS, to construct a recombinant plasmid, pET30a-GnRH5-MT3, which was expressed and purified using E. coli BL21(DE3) cells. The recombinant protein was best expressed under the induction conditions of 25 ℃ with 0.25 mM IPTG, with a purity of up to 80% after purification. It was mixed with the external adjuvant ISA 206 and immunized mice by subcutaneous injection. Serum levels of GnRH-specific Antibody, T, FSH and LH were detected by ELISA, and testicular tissue morphology was observed by HE staining. The results showed that GnRH5-MT3 recombinant protein combined with external adjuvant could efficiently induce GnRH-specific antibody production in male mice (*P* < 0.01–0.001) and significantly reduce the Serum LH, FSH and T concentrations (*P* < 0.01–0.001). The immunized mice had sparse cells in the seminiferous tubules of the testis, significantly reduced cells at various developmental stages, and almost no mature spermatozoa. This study provides experimental basis for further application of the vaccine in animal immune castration.

## Introduction

In the field of animal production and veterinary medicine, the regulation of gonadal function is of paramount importance. Castration, as a common management method, primarily uses surgical or chemical agents to remove or functionally inhibit the reproductive glands (testes or ovaries) of animals to eliminate their reproductive capacity and reduce sex hormone levels ^[1-2]^. However, the method has obvious limitations, and immune castration has gradually gained attention as an alternative modality due to considerations of animal welfare ethics and the productive benefits of animal husbandry. Immune castration refers to the suppressing of the secretion of gonadal hormones and the development of the sexual organs by using immune means to achieve castration effect _[3–4]_.The core mechanism is to activate the body’s immune system, induce the production of high titer specific antibodies, neutralize endogenous key hormones, and block relevant regulatory pathways, thereby achieving the goal of controlling fertility ^[5–6]^.Compared with traditional castration methods, this method has the characteristics of precise operation, reversible effect and few side effects, and has broad application prospects.

Gonadotropin-releasing hormone (GnRH) is a decapeptide hormone secreted by the hypo thalamus and plays a central regulatory role in the reproductive system of mammals, integrating and regulating reproductive processes and related behaviors mainly through the hypothalamic-pituitary-gonadal (HPG) axis ^[7-8]^.Studies have shown that the application of a GnRH immune strategy can induce the body to produce specific antibodie that neutralize endogenous GnRH, thereby blocking the reception of pituitary signals, inhibiting downstream reproductive function, and disrupting the inherent balance of the body’s HPG axis, thus achieving the purpose of immune castration ^[9-10]^. However, due to the weak molecular immunogenicity of GnRH, it is often conjugated with carrier proteins or combined with external adjuvants in practical vaccine development to enhance its ability to elicit an immune response. Research by Pan F et al. demonstrated that a two-copy GnRH tandem construct combined with ISA 201 adjuvant could induce high-titer antibodies and effective castration in animal models ^[11]^.

Metallothionein-3 (MT3) is a metal-binding protein with a molecular weight of approximately 7 kDa, consisting of two domains, the α and β domains which provide four and three zinc-binding sites respectively. Its adjuvant effect relies on fusion with antigens and regulation of zinc homeostasis ^[12-15]^. Zinc ion balance is crucial for the proper function of the immune system ^[16]^. Studies indicate that after exogenous MT3 enters the body, a large number of free zinc ions are captured by thiol groups, forming stable complexes in the neutral extracellular environment. After fused with antigens, the fusion protein can directly activate dendritic cells (DCs), promote germinal center formation, and accelerate antibody class switching. YIN et al. for the first time, used human MT3 as a molecular adjuvant fused with Omp19 and HC, respectively, showing that even with reduced antigen doses, greater immunogenicity can be induced, and the effect is better than mixed use of traditional external adjuvants ^[17]^. This suggests that human MT3 can promote rapid, highly efficient, and sustained antibody responses to protein antigens, demonstrating its potential as a novel built-in adjuvant for vaccine design.

Traditional Immune castration vaccines often rely on exogenous adjuvants to non-specifically stimulate the immune system ^[18-20]^. However, this strategy has problems such as uneven response intensity, significant individual variability, multiple booster immunizations required, and potential side effects ^[21-22]^. Therefore, the combination of exogenous adjuvants with endogenous “adjuvant signal” molecules is expected to achieve synergistic enhancement of immune responses.

Although the strategy of combining endogenous molecular adjuvants with external adjuvants has been reported in some vaccine studies, the current application of this strategy in the field of castration vaccines has not been systematically elucidated in the literature. This study aims to evaluate the immunogenicity of a fusion protein comprising MT3 and a GnRH pentamer combined with the external adjuvant ISA 206 through animal experiments, thereby exploring the impact of this strategy on the efficacy of GnRH-based Immune castration vaccines.

## Materials and methods

### Plasmids, bacterial strains, peptides, and experimental animals

The pET-30a(+) plasmid was synthesized by GenScript Biotech Corporation (Nanjing, China). The E. coli BL21(DE3) strain was purchased from TransGen Biotech (Beijing, China) and preserved in our laboratory. The GnRH pentamer peptide was synthesized by GL Biochem (Shanghai) Ltd. Thirty 4-week-old healthy male Balb/c mice were purchased from Changchun Yisi Experimental Animal Technology Co., Ltd. The mice were housed under the following conditions: clean-grade animal room (barrier system) at Changchun Xinuo Co., Ltd., with room temperature maintained at 19–23°C, relative humidity at 40–65%, alternating light 12 h bright and 12 h dark. Mice were kept in individually ventilated cages (IVCs) with free access to autoclaved food and water.

All animal experiments were approved by the Animal Ethics Committee of Changchun Longsheng Experimental Animal Technology Co., Ltd. (CCLSLL-2024110703) and the experimental operations were subject to the “Guidelines for the Welfare and Ethics of Experimental Animals in China”.

### Construction and expression of recombinant plasmid

The Human-origin MT3 sequences (NM_005945.4) and GnRH sequences (PO7490) were searched on NCBI. MT3 was linked to the N- and C-termini of the GnRH pentamer by the flexible linker GGGGS, with a 6×His tag added to the C-terminus, as shown in Fig. 1, and the gene sequence was optimized for E. coli adapted codon optimization. Two enzymatic cleavage sites, BamH I and Nde I, were selected. The optimized sequence was cloned into the pET-30a() vector (synthesized by Nanjing Jinsrui Biotechnology Co., Ltd.), the plasmid pET30a-GnRH5-MT3 was constructed, and sequencing identification was performed. Recombinant plasmid pET30a-GnRH5-MT3 was converted into E.coli BL21(DE3) competent cells, cultured overnight at 37 °C at 180 rpm in LB medium containing 50 μg/mL kanamycin, and when OD values reached 0.6-0.8, IPTG was added to a final concentration of 0.5 mmol/L, and expression was induced at 25 °C at 220 rpm for 12 h.

**Figure 1.**
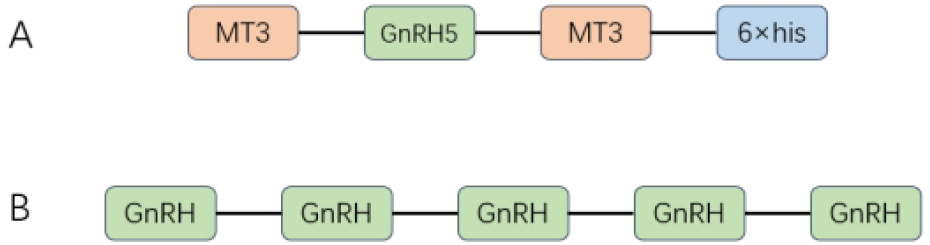
Schematic illustration of the GnRH5-MT3 and GnRH5-based antigen constructs.

### Purification of the recombinant protein

The Postinduction bacterial juice was collected at 4°C with centrifugation at 12000 r/min for 12 min, the bacteria were resuspended with PBS, and ultrasound was performed at 320 W in an ice-bath environment for 25 min; protein expression was analyzed by SDS-PAGE for pre-induction, post-induction, ultrasound supernatant, and ultrasound precipitation. 35mL of ultrasound supernatant was taken, sterilized by filter with a pore diameter of 0.45 μm, purified by Ni column, eluted with different concentrations of imidazole, and purified samples were obtained. After SDS-PAGE identification, imidazol was removed by changing the solution in a dialysis bag of 5KD, placed in 1L of PBS solution (pH=7.4) for three dialysis sessions, and changed every 8h.Protein concentrations were accurately determined by BCA Protein Assay kit by centrifugation of dialyzed protein samples at 4 °C, 5000 rpm for 20 min using a 5kD ultrafilter tube.

### SDS-PAGE analysis of recombinant protein

Protein samples were pre-prepared with a 12% polyacrylamide gel, mixed with a 4 × Loading Buffer, boiled in boiling water for 10 min, and after adding the samples, 80V voltage was run to concentrate the gel and 120V voltage was run to separate the gel. After the electrophoresis, the separate gel was cut and placed in the staining solution for staining, and after 30 min, the staining solution was used for decolorization until the separation gel background was clear.

### Animal immunization protocol

Four-week-old male Balb/c mice were selected and placed in greenhouse in 12-h light and 12-h dark cycles with free feed and drinking water. Thirty mice were randomly divided into five groups, each with the first immunization at 0 wk, booster immunization at 2 wk and 4 wk, Serum was collected every 2 wk, and mice were sacrificed at 10 wk, as shown in Figure 2. The immunization was done by subcutaneous injection with protein immunization groups GnRH5, GnRH5 MT3, GnRH5 ISA 206, GnRH5 MT3 ISA 206 and PBS control arm. The recombinant protein dose of the immunized group was 30 μg per mouse, in which ISA 206 adjuvants were mixed with recombinant protein in a 1:1 ratio to prepare the vaccine. PBS was diluted to 100 μL as shown in Table 1.

**Figure 2.**
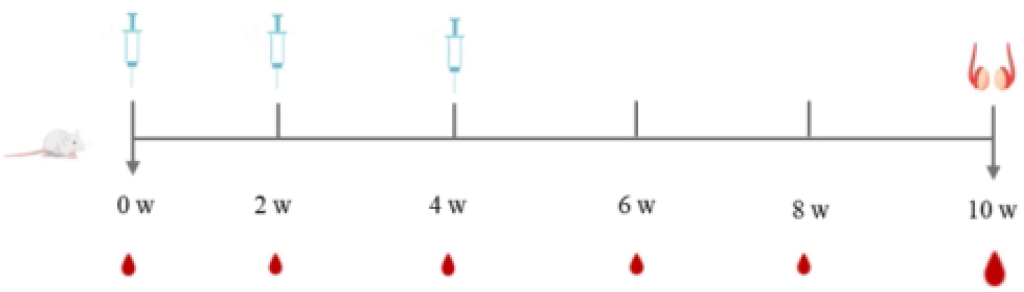
Schematic diagram of the immunization schedule.

**Table 1.**
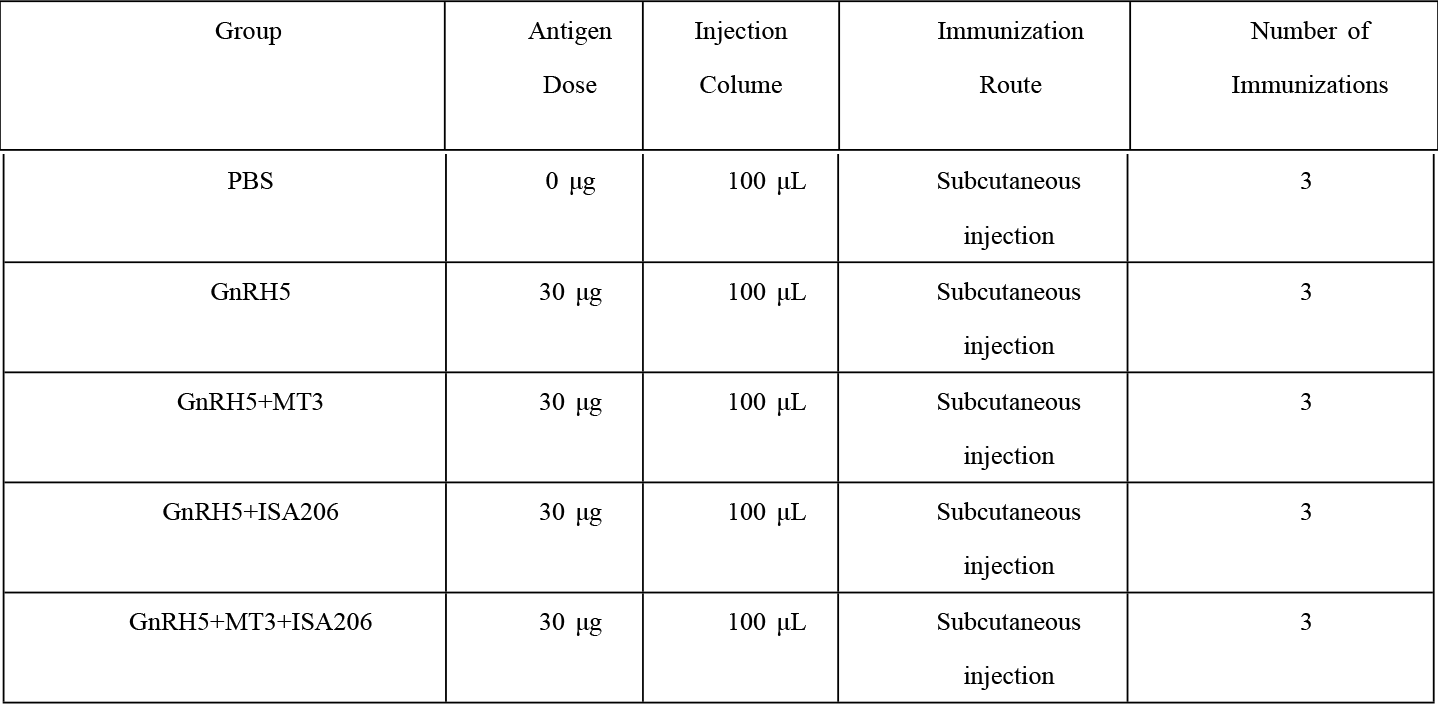
Experimental grouping and immunization dosage.

The experimental animals and experimental operations involved in this article have been approved by the Ethics Committee and strictly follow the rules and regulations of animal experiments. After the experiment, all experimental animals were treated humanely (amniotic fluid) to minimize their pain and harm, in accordance with animal ethics related norms.

### Sample collection

Peripheral blood samples were collected 150 μL every 2 w from the tail vein of each mouse, coagulated in a centrifuge tube for 1 h, centrifuged at 3000 rpm for 30 min, Serum was collected and stored at -80 °C. Mice were sacrificed and testis were removed at 10 w postimmunization and stored in 4% paraformaldehyde solution. (Beijing Reagan Biotechnology Co. CM0052).

### Detection of serum anti-GnRH antibodies by ELISA

Flat-bottom 96-well plates were coated with 1 μg of GnRH polypeptide in coating buffer (100 μL/well, pH 7.4) and incubated overnight at 4 °C. Plates were washed 5 times with 0. 05% (v/v) Tween 20 for 1 min, followed by 2% bovine Serum Albumin, ALB (200 μL/well) for 1 h at 37 °C. Discard the blocking buffer, wash 5 times with PBS buffer containing 0.0 5% (v/v) Tween 20, add dilute Serum (200 μL/well; 1:50) to the sample in duplicate and block at 37 °C for 1 h. After washing 5 times with PBS buffer containing 0.05% (v/v) Tween 20, HRP-conjugated goat anti-mouse IgG Antibody 1:5000 (Beyotime, China) was added, the plates were incubated for 1 h at 37 °C, the plates were washed 5 times, and 100 μL TMB Substrate was added to each well for 10 min of light-avoidance reaction. The reaction was stopped by adding 50 μL 2 M sulfuric acid solution to each well. The absorbance of each well was measured with a microplate reader at a wavelength of 450 nm, and the results are expressed as the optical density (OD) value

### Measurement of serum testosterone (T), follicle-stimulating hormone (FSH), and luteinizing hormone (LH) levels

The concentrations of T, FSH, and LH in serum were measured using commercialenzyme-linked immunosorbent assay (ELISA) kits according to the manufacturers’ instructions: Mouse T ELISA Kit (Wuhan Saier Biotechnology Co., Ltd., Cat: SF14088) ; Mouse FSH ELIS A Kit (Wuhan Saier Biotechnology Co., Ltd., Cat: SF14079) ; Mouse LH ELISA Kit (Wuhan Saier Biotechnology Co., Ltd., Cat: SF14102)

### Hematoxylin and Eosin (H&E) staining of testicular tissues

After euthanasia, testes were carefully dissected, and surrounding blood vessels, adipose tissue, and connective tissues were removed. The tissues were then processed for routine H& E staining.

### Statistical analysis

All statistical analyses were performed using GraphPad Prism software. Statistical comparisons between different study groups were performed with t tests and one-factor analysis of variance. In all tests, statistical significance was marked as follows: ^*^*P*<0.05,^**^*P*<0.01,^***^*P*<0. 001

## Results

### Restriction enzyme digestion analysis of therRecombinant plasmid pET30a-GnRH5-MT3

The double-enzymatic identification of the recombinant plasmid pET30a-GnRH5-MT3 was performed by selecting the NdeI, BamHI restriction site for enzyme cleavage. The results of 1% agarose gel electrophoresis are shown in Figure 3. The enzyme cleavage product exhibits two specific bands in the electrophoresis spectrum, which are consistent with the theoretical predicted values of the empty plasmid pET-30a and GnRH5-MT3 fusion genes, respectively, indicating that the recombinant expression plasmid has been accurately constructed.

**Figure 3.**
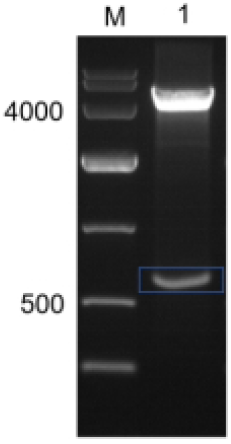
Results of double enzymatic isolation of recombinant expression plasmid.: Lane M: DL10,000 DNA Marker; Lane 1: Recombinant plasmid pET30a-GnRH5-MT3 digested with Nde I and BamH I.

### Expression of the Recombinant Protein

The recombinant plasmid pET30a-GnRH5-MT3 was transformed into E. coli BL21(DE3) competent cells and induced to express by 0.25 mM IPTG at 25 °C. The bacterial juice was analyzed by SDS-PAGE after ultrasound fragmentation. The results are shown in Figure 4. Specific bands of approximately 23 kDa were seen in lanes 2, 3, and 4, with molecular mass sizes consistent with GnRH5-MT3 proteins, indicating that the SF protein was successfully expressed and that most of the protein was present in soluble form.

**Figure 4.**
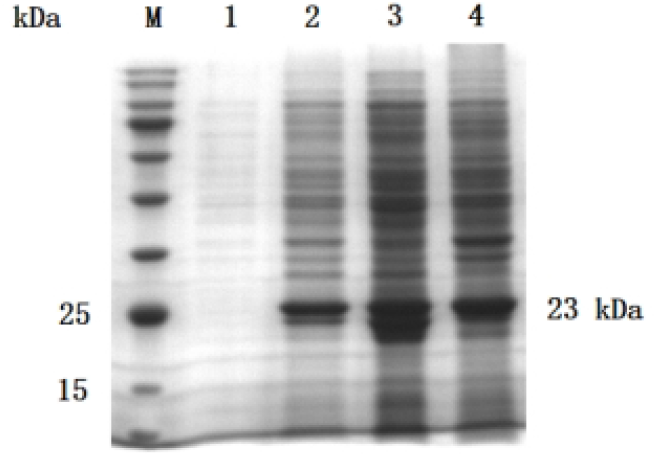
Analysis of recombinant protein expression. M: Protein Molecular Weight Marker; Lane 1: pET-GnRH5-MT3 before induction; Lane 2: pET-GnRH5-MT3 after induction; Lane 3: Soluble fraction (supernatant) after ultrasonication of pET-GnRH5-MT3; Lane 4: Insoluble fraction (pellet) after ultrasonication of pET-GnRH5-MT3.

### Purification of the recombinant protein

After affinity chromatography purification, the recombinant protein was further purified using a NanoGel-50Q ion exchange column. As shown in Fig. 5, the protein purified under this condition exhibited the highest purity. Therefore, this method was selected for subsequent experimental use. The concentration of the purified protein, as determined by the BCA assay, was 1.25 mg/mL.

**Figure 5.**
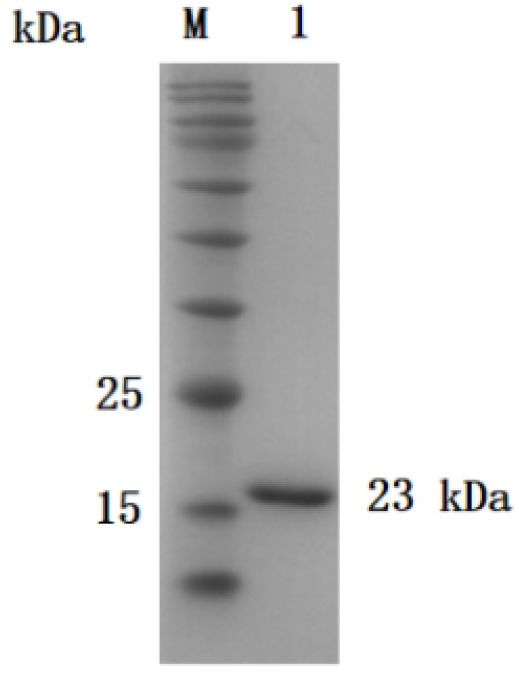
Purification profile of the recombinant protein. M: Protein Molecular Weight Marker; Lane 1: Purified GnRH5-MT3 fusion protein.

**Figure 6.**
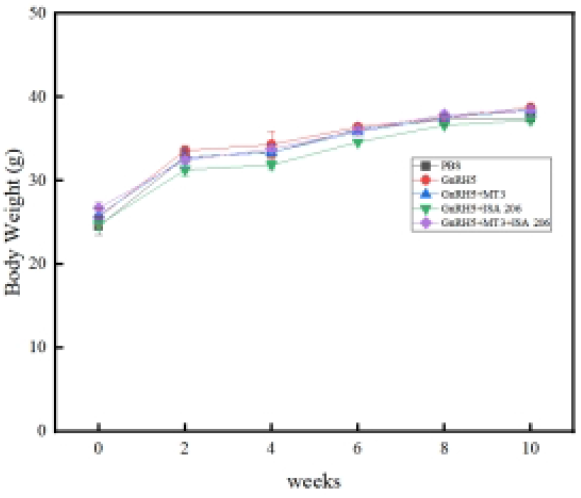
Body weight changes in mice following immunization.

### Body weight monitoring during immunization

Body weights of immunized mice were monitored throughout the study. As shown in Fig. 9, no significant changes in body weight were observed in any of the immunized groups compared to the PBS control group following vaccine administration. All groups exhibited a steady increase in body weight over time, with no statistically significant differences compared to the control group (*P* > 0.05).

### Specific antibody titers in mice serum

Anti-GnRH specific antibody levels in mouse serum were measured by ELISA. As shown in Fig. 7, all immunized groups developed GnRH-specific antibodies after the second immunization. The specific antibody levels increased gradually, reaching a peak at week 8. This immune response persisted through week 10. No anti-GnRH antibodies were detected in the control group.

**Figure 7.**
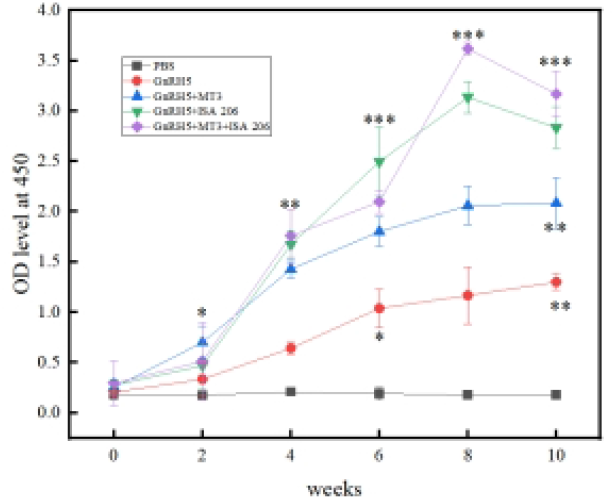
Determination of GnRH-specific antibody levels in serum of vaccinated groups and control groups. ^*^*P* < 0.05, ^**^*P* < 0.01, ^***^*P* < 0.001 (t-test).

### Determination of T, FSH and LH concentrations in serum

T concentrations in Serum were detected every two weeks by ELISA kits. As shown in Figure 8a, T levels in each immune group decreased gradually after immunization and remained significantly below the concentration level of control arm and remained until ten weeks after the first immunization.

**Figure 8.**
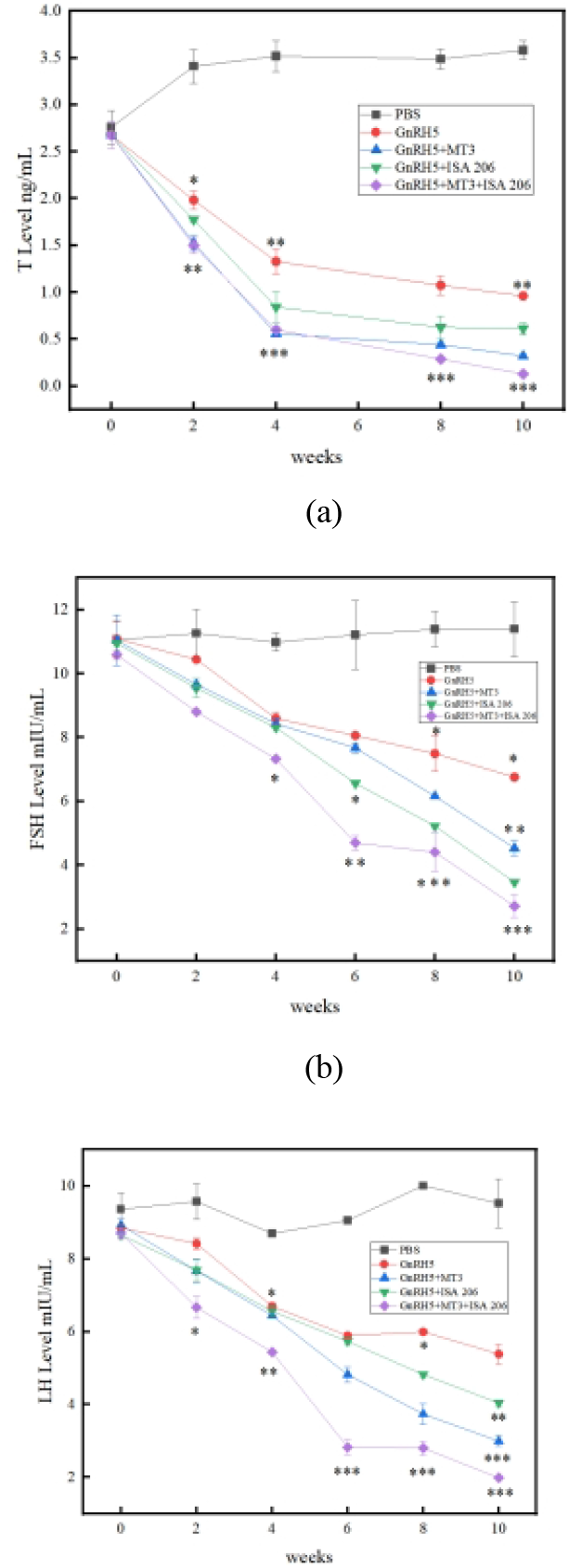
(a) Determination of T content in Serum (ng/mL). (b) Determination of FSH in Serum (mIU / mL). (c) Determination of LH content in Serum (mIU/mL) ^*^*P* < 0.05, ^**^*P* < 0.01, ^***^*P* < 0.001 (t-test).

FSH and LH concentrations in Serum were detected every two weeks. As shown in Figures 8b and c, Serum FSH and LH concentrations in the immunized group decreased gradually two weeks after the first immunization, reaching their lowest level at ten weeks after the immunization, significantly lower than in the blank control arm.

### Histological analysis of testes by H&E staining

At the 10th week, mice were euthanized, and testes were collected for histological examination via hematoxylin and eosin (H&E) staining.

As shown in Fig. 9a, testicular tissues from the PBS control group exhibited normal seminiferous tubule morphology, with abundant spermatogenic cells at various developmental stages—including spermatogonia, primary spermatocytes, secondary spermatocytes, spermatids, and mature spermatozoa—closely arranged within the tubule lumina.

**Figure 9.**
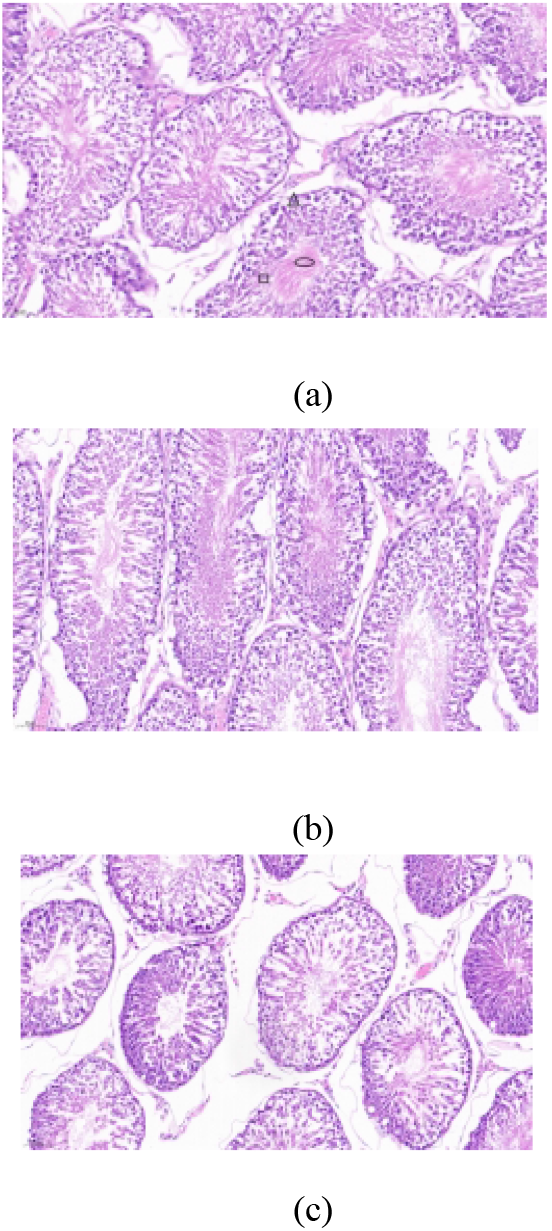
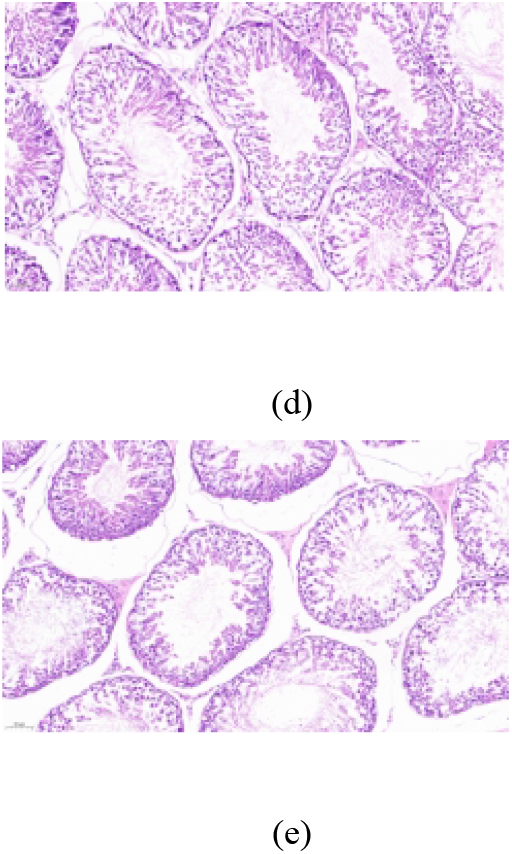
Histological sections of testicular tissue with H&E staining. (a) Testis from PBS control group. Box: spermatogonia; Triangle: spermatids; Circle: spermatozoa. (b) Testis from GnRH5 immunized group. (c) Testis from GnRH5+MT3 immunized group. (d) Testis from GnRH5+ISA 206 immunized group. (e) Testis from GnRH5+MT3+ISA 206 immunized group.

In contrast, tissues from the GnRH5 (Fig. 9b), GnRH5+MT3 (Fig. 9c), and GnRH5+ISA 206 (Fig. 9d) vaccine groups showed reduced cellularity within the tubules, inhibition of spermatid development, and a marked decrease in sperm count.

Most notably, the GnRH5+MT3+ISA 206 vaccine group (Fig. 9e) displayed severely sparse cellularity within the lumina, with very few cells at any developmental stage and an almost complete absence of sperm.

## Discussion

As an animal-friendly castration technique, immuno castration is centered on the challenge of a highly efficient immune response by enhancing the immunogenicity of GnRH ^[23-25]^. In this study, five GnRH epitopes were protein recombined with the built-in adjuvant MT3 using genetic engineering techniques, which efficiently initiated the immune response and improved immunogenicity by providing high-density repeat epitopes. Recombinant proteins have tension during expression, and the use of a flexible linker of GGGGS ensures independent folding and spatial synergy of antigen with adjuvant ^[26-27]^, which lays a structural basis for efficient initiation of immune responses. The soluble expression and successful purification of recombinant protein (with purity up to 80 %) provide a stable material basis for subsequent animal experiments.

GnRH-specific Antibodies refer to immunoglobulins produced by the mouse immune system under stimulation with GnRH antigens that are capable of highly specifically recognizing and binding to GnRH ^[28-31]^. In this study, the response effects of different immune groups were assessed by regular (two-weekly) measurements of the antibody levels in the mouse serum. Both GnRH5-specific antibodies were successfully induced in mice after immunization with GnRH5 and GnRH5+MT3, but the GnRH5+MT3 group showed higher antibody titers than the GnRH5 group, suggesting that the use of the built-in adjuvant MT3 induced a more efficient immune response.

LH promotes the synthesis and secretion of T by acting on the Leydig cells of the testis, and FSH, by acting on the seminiferous tubules of the testis, synergizes with T to initiate and maintain the process of sperm production, both of which synergize to maintain normal physiological functions of the testis ^[32–34]^. The results of this study showed that LH, FSH, and T concentrations in mice decreased to varying degrees after vaccination with GnRH5 and GnR H5 MT3, and the decrease was more significant in the GnRH5 MT3 group than in the GnR H5 group. This might be due to the fact that the higher levels of GnRH antibodies in this group neutralized the endogenous GnRH in mice, resulting in ineffective absorption of GnRH signaling by the pituitary gland, which in turn inhibited the release of FSH and LH, and ultimately inhibited the synthesis and secretion of downstream T.

Internal molecular adjuvants function through specific molecular pathways targeting the Immune System, demonstrating significant advantages in areas such as vaccine development and immunotherapy ^[35–36]^. Halbroth BR et al., fusion expression of the invariant chain of MHC class II molecules (Ii) with target antigen genes as molecular adjuvants, showed that the fusion protein was able to significantly enhance antigen-specific CD4 and CD8 T cell immune responses ^[37]^. This genetically engineered technique allows direct fusion of adjuvant-active molecules with target protein antigens to construct a single, self-adjuvanted recombinant molecule that not only achieves precise delivery of antigens, but also effectively activates the Immune System and optimizes the quality and persistence of immune responses.

In vaccine development, the choice of an external adjuvant is one of the key strategies to enhance the intensity and durability of the immunogenic induced immune response ^[38-40]^. As an optimized oil-emulsion adjuvant, ISA 206 has its unique water-in-oil-in-water (W/O/W) structure that efficiently encapsulates the antigen and achieves slow release, thereby inducing the body to produce high levels and durable neutralizing Antibodies, ultimately triggering a stronger and more durable immune response ^[41-43]^.

Based on the above characteristics, the recombinant protein was used in combination with the external adjuvant ISA 206 in this study. The experimental results showed that the immune group used in combination with internal and external adjuvants was superior to the immune group used with either internal or external adjuvants alone on several key indicators, including specific antibody levels, and concentrations of T, FSH, and LH. By means of P-value analysis of the data, antibody levels increased in all immunization groups 2 weeks after the first immunization, and were significantly higher in the combined intra- and extra-adjuvant doses than in the monoadjuvant doses; meanwhile, FSH, LH, and T concentrations decreased significantly after 2 weeks of immunization, with T concentrations dropping below 1 ng/mL after 4 weeks of immunization and remaining low until the 10th week. Because the development and function of the testis depend on the secretion of T cells ^[44–47]^, we further performed morphological analysis of the testicular tissue in mice. The control arm had normal structures of the seminiferous tubules and normally developed and differentiated germ cells visible in the lumen, whereas in the GnRH5+MT3+ISA 206 combined with internal and external adjuvants, almost no mature spermatozoa were seen in the testicular lumen, and the number and quality of germ cells were significantly lower in the GnRH5+MT3 alone-contained adjuvant group. The above results suggest that our vaccine, by continuously maintaining high levels of GnRH Antibody, disrupts the physiological balance of the HPG axis, effectively reduces T concentration, thereby inhibiting the development and function of the testes and ultimately inducing irreversible atrophy.

Safety assessments showed that the experimental group of mice showed no significant difference in body weight growth compared with control arm (P>0.05), indicating that the vaccine and adjuvant combination had no significant systemic toxicity, demonstrating that the GnRH recombinant vaccine and its adjuvant combination were well established in the immunization process.

In summary, our study constructed a recombinant GnRH vaccine with MT3 protein as an internal adjuvant and combined with an external adjuvant, ISA 206, to explore its immune response characteristics in mice. Compared with conventional vaccine formulations using only an internal adjuvant or an external adjuvant, it induced higher levels of GnRH-specific antibodies in the body, achieving long-term hormone inhibition and a significant decrease in sperm quality, significantly improving castration effects, which is of great guiding significance for the design and optimization of vaccine adjuvants. Future exploration of the immune validity period of the vaccine is also needed, while exploration of the composite use of internal and external adjuvants or the use of other types of adjuvants.

## Acknowledgments

We would like to extend our sincere gratitude to Changchun Sinor Co., Ltd. Additionally, we appreciate Liu Bing from Jilin Wuxing Co., Ltd., for his help in serum sample processing.

## Author contributions statement

Data curation, G.Z., Y.X., and Y.Y. ; Formal analysis, Y.X., G.Z. and Y.Z.; Funding acquisition, Y.Y.; Investigation, Y.Z.; Methodology, Y.X. and G.Z.; Project administration, Y.Y.; Resources, Y.Y.; Software, Y.X.; Supervision, C.W. and Y.Y.; Validation, Y.Y. and C.W.; Visualization, Y.X. and G.Z.; Writing—original draft, G.Z and Y.X; Writing—review and editing, C.W. and Y.Y. All authors have read and agreed to the published version of the manuscript.

## Additional information

### Institutional review board statement

The animal study protocol was approved by the Animal Ethics Committee of Changchun Long Sheng Experimental Animal Technology Co., Ltd. (Approval Code: CCLSLL-2024110 703; Date: 7 November 2024), and the experimental operations were subject to the “Guidelines for the Welfare and Ethics of Experimental Animals in China”.

### Data availability statement

The data presented in this study are contained within the article.

